# Epitranscriptome analysis of NAD-capped RNA by spike-in-based normalization

**DOI:** 10.1101/2023.03.23.534034

**Authors:** Dean Li, Shuwen Ge, Yandong Liu, Miaomiao Pan, Xueting Wang, Guojing Han, Sili Zou, Rui Liu, Kongyan Niu, Chao Zhao, Nan Liu, Lefeng Qu

## Abstract

The hub metabolite, nicotinamide adenine dinucleotide (NAD), can be used as an initiating nucleotide in RNA transcription to result in NAD-capped RNA (NAD-RNA). NAD-RNA that intimately connects metabolite with gene expression can be developed as novel biomarkers for aging and disease. Epitranscriptome-wide profiling of NAD-RNAs involves chemo-enzymatic labeling and affinity-based enrichment; yet currently available computational analysis cannot adequately remove variations associated with capture procedures. Here, we propose a spike-in-based normalization and data-driven evaluation framework, enONE, for the omic-level analysis of NAD-capped RNAs. We demonstrate that carefully designed spike-in RNAs, together with modular normalization procedures and evaluation metrics, can lead to the optimal normalization that maximally removes unwanted variations, empowering quantitative and comparative assessment of NAD-RNAs from different datasets. Using enONE and a human aging cohort, we reveal critical features of NAD-capped RNAs that occur with normal age. enONE facilitates the discovery of NAD-capped RNAs that are responsive to physiological changes, laying a critical foundation for functional investigations into this modification.

## INTRODUCTION

NAD, an adenine nucleotide containing metabolite, can be incorporated into the RNA 5’-terminus to result in NAD-capped RNA (NAD-RNA)^1,2^, which is different from eukaryotic canonical cap structure predominantly formed by 7-methylguanosine (m^7^G) via a 5’-to-5’-triphosphate bridge (m^7^G-RNA)^3,4^. It has been estimated that NAD-capped forms make up more than 0.6% and 1.3% of the genes expressed in the entire transcriptome from mouse liver and kidney, respectively^5^. To capture such low-level capping events, the recently developed NAD-RNA identification methods involve the use of chemo-enzymatic reaction, followed by affinity-based enrichment^6-8^. However, the resulting high-throughput sequencing data can be hampered by the effect of capture procedures and other unwanted variations. Given these limitations, current computational methods cannot be directly applied to the omic-level assessment of NAD-capped RNAs.

Normalization is an essential step to remove unwanted variations. N^6^-methyladenosine (m^6^A), a prevalent epitranscriptomic modification in RNA, has been extensively characterized in virus and eukaryotic organisms^9^. Computational tools for m^6^A-seq, e.g., RADAR^10^ and m^6^A-express^11^, employ a split scaling strategy that calculates scale factors for input and enrichment, respectively, to adjust the variations from enrichment procedures and sequencing depth. However, these analytical methods cannot properly account for the unwanted variation between samples with and without enrichment, thus challenging the identification of enrichment signals. More generally, current analyses of epitranscriptomic data are mostly based on normalization designed for bulk RNA-seq, e.g., scaling-based methods, such as Total Count (TC), Trimmed Mean of M values (TMM)^12^, and DESeq^13^. The implicit assumption underlying scaling-based methods is that all the gene-level counts are proportional to scale factors and that the between-sample variations can be adequately adjusted by scale factors. Unfortunately, this assumption is inevitably violated when affinity-based enrichment selectively amplifies the signal of genes, e.g., m^6^A and NAD-capping, which leads to disproportional gene counts between input and enrichment. Another regression-based method, Remove Unwanted Variation (RUV) ^14^, regress gene count measurements on unwanted factors, thus computing corrected expression values from the residuals. The implicit assumption underlying this method is that a set of negative controls, which are not affected by covariates of interest, is available, such as the spike-in from the External RNA Controls Consortium (ERCC). However, the ERCC-based method suffers from discrepancies between endogenous transcripts and spike-in, hindering its usage in omic-level profiling. Limited by current analytical methods, nuisance variations fail to be properly corrected for NAD-RNA sequencing data, thereby obscuring true biological signals.

NAD-modified RNA connects metabolite with gene expression. NAD is the hub metabolite and redox regent for cells, involving in a wide range of biological processes^15^. In rodents and humans, studies have revealed that the NAD level declines with age in critical tissues and organs^16^. Given the dynamics of NAD and gene expression over the course of adult lifespan, NAD-capped RNAs, poised to integrate metabolomics and transcriptomics, may provide novel insights into physiological and perhaps pathological situations. Thus, it is tempting to explore how NAD-modified epitranscriptome is modulated with age. In the present study, we develop enONE framework for NAD-capped RNA analysis by spike-in-based omic-level normalization and evaluation. enONE integrates spike-in controls, global scaling, and regression-based normalizations, followed by performance evaluation that selects the local-optimal normalization method to remove unwanted variation. Using human aging cohort, we apply enONE to the identification of NAD-RNAs from circulating blood cells, revealing dynamics of NAD modification with age.

## RESULTS

### The workflow of enONE

Using exogenous spike-in RNAs, we designed a computational framework that integrates global scaling and regression-based normalization modules. We thereby named our analytical method enONE, for **<u>E</u>**pitranscriptional **<u>N</u>**AD-capped RNA analysis by spike-in-based **<u>O</u>**mic-level **<u>N</u>**ormalization and **<u>E</u>**valuation (**Fig. 1**).

**Figure 1:**
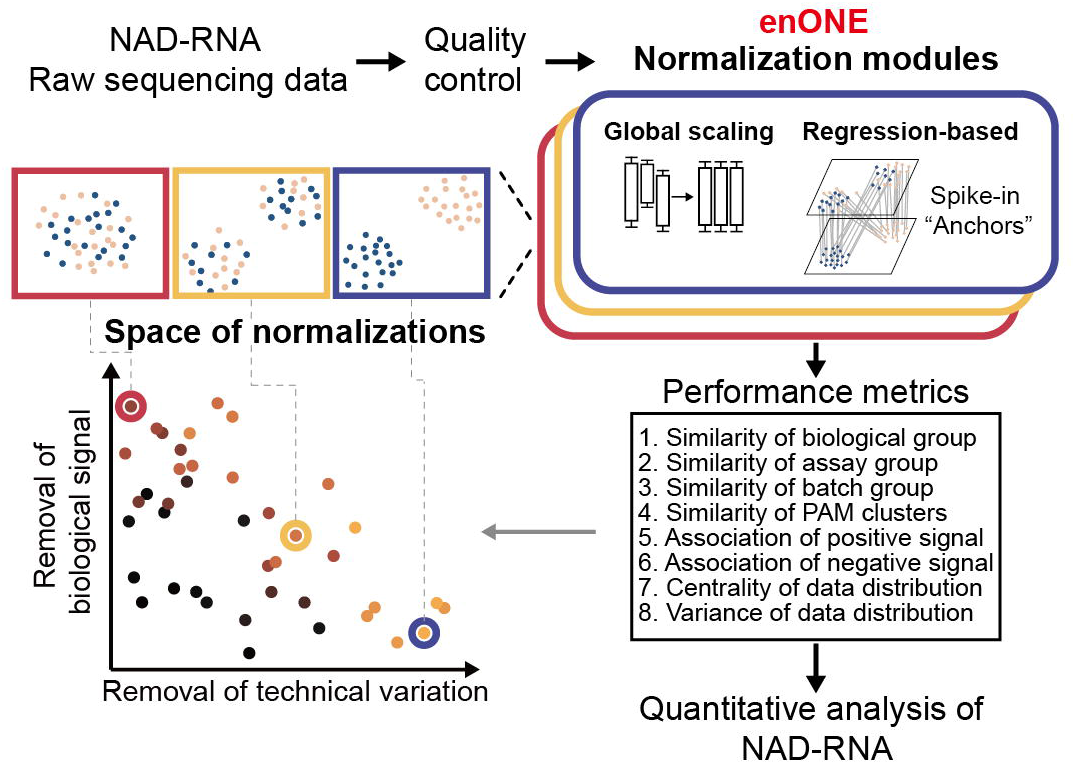
The workflow of enONE. enONE starts with quality control to obtain high-quality sequence reads by removing outliers and lowly-expressed genes. enONE performs spike-in-based normalization that integrates global scaling and regression-based procedures to generate normalization toolsets. Next, enONE uses eight data-driven metrics to evaluate normalization performance. By exploiting the full space of normalizations, enONE identifies the optimal procedure that maximally removes unwanted variations, while minimally impacting the signals from NAD-RNA-seq data.

enONE initiates with a quality control step, to remove outlier samples and low-expressed transcripts. Second, a subset of genes from spike-in RNAs, with their expressions presumably not being influenced by the covariates of interest (e.g., enrichment assay or biological condition), are used as the anchor set to estimate unwanted variation (e.g., batch effect). A generalized linear model (GLM) is then applied to regress the observed read counts from anchor set on the unknown nuisance variables to estimate factors that are subsequently used by the normalization tools for the adjustment of unwanted variation. Third, a two-part normalization template is employed to define an ensemble of the normalization procedures: 1) global scaling of read counts to account for between-sample difference in sequencing depth and other parameters of the read count distribution, and 2) regression-based adjustment for unwanted variations. For instance, one can apply a robust scaling procedure, such as TMM, followed by unsupervised procedures to estimate hidden unwanted variations and regress them out of the data (e.g., RUV^14^). Fourth, enONE comparatively analyzes all normalization toolsets to identify sets of top-performing procedures. Specifically, enONE calculates ranks based on eight performance metrics that represent the local-optimal trade-offs towards removing unwanted variation, preserving biological variation of interest, and maintaining minimum technical variability of global expression. Combined, enONE utilizes a data-driven approach to determine appropriate normalization procedures for the quantitative analysis of NAD-modified epitranscriptome.

### Epitranscriptomic profiling of human PBMCs

To gain insights of NAD-modified epitranscriptome during aging, we collected human peripheral blood mononuclear cells (PBMCs) from an aging cohort in community subjects comprising of young (N = 23, age: 23-32), middle (N = 20, age: 40-50), and old (N = 18, age: 54-67) individuals for epitranscriptome-wide profiling of NAD-RNAs (**Fig. 2A**), according to the inclusion criteria approved by the Ethics Committee. Clinical characteristics of the participants were evaluated and listed in Supplementary Table 1. As an essential component of enONE, we deliberately included three types of spike-in RNAs: 1) total RNAs from *Drosophila melanogaster*, an invertebrate model organism with well-annotated genome sequence; 2) synthetic RNAs, consisting of 5% NAD-relative to m^7^G-capped forms, were used to determine the capture sensitivity; 3) synthetic RNAs, with 100% m^7^G-capped forms, were used to determine the capture specificity (**Fig. 2A**). Notably, spike-ins 2 and 3 were synthesized with templates from different sequences. Combined, we subjected 10 μg total RNAs from human PBMCs mixed with 40 μg total RNAs from *Drosophila*, 0.1 ng spike-in 2 RNAs, and 0.1 ng spike-in 3 RNAs to NAD-RNA sequencing, followed by enONE computational analysis of NAD-RNA profiles.

**Figure 2:**
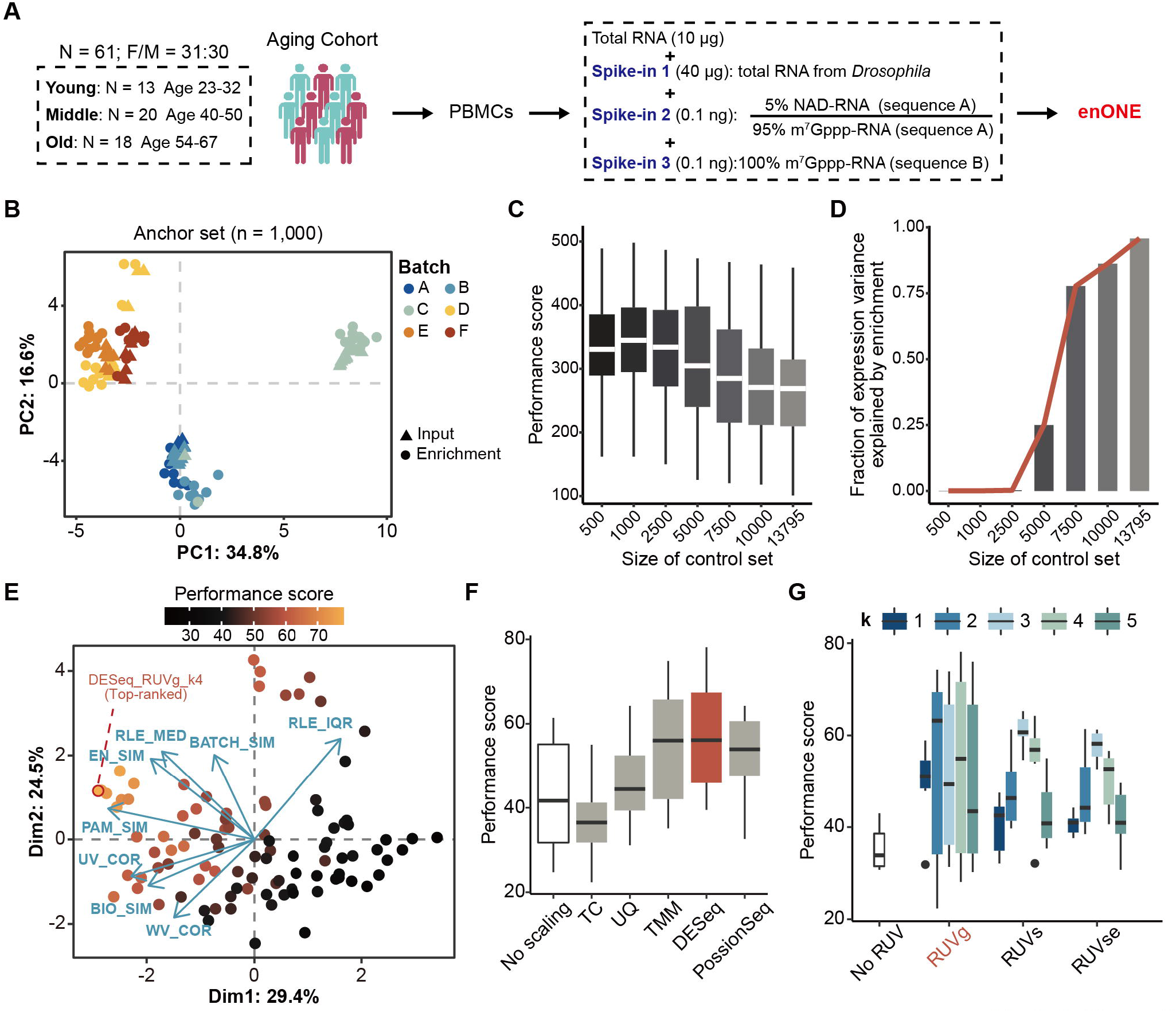
The feasibility of enONE. (A) Aging Cohort and experiment outline. A total of 61 participants (female/male = 31:30, age 23-67) were enrolled for NAD-modified epitranscriptome profiling. Total RNAs from PBMCs were mixed with *Drosophila* spike-in RNA, and two synthetic spike-in, of which one with 5% NAD-capped forms and another with 100% m^7^G-capped forms. The mixture was subjected to NAD-RNA-seq, followed by enONE computational analysis of NAD-RNA profiles (highlighted in red). (B) PCA of the 1,000 least significantly enriched genes (denoted as anchor set) from *Drosophila* spike-in counts. (C) Boxplot showing the normalization performance based on anchor set of different sizes from *Drosophila* spike-ins. (D) Fraction of the first two expression PCs variance explained values (taken cumulatively) for linear regression model using enrichment variable as the explanatory variable. PCs are computed from the variance stabilizing and transformed matrix of spike-in counts. (E) enONE identified the top-ranked procedure (DESeq_RUVg_k4) from a total of 96 procedures. Each point corresponds to a normalization procedure and is colored by the performance score (mean of eight scone performance metric ranks). The blue arrows correspond to the PCA loadings for the performance metrics. The direction and length of a blue arrow can be interpreted as a measure of how much each metric contributed to the first two PCs. (F) Boxplot showing performance score, stratified by scaling procedures. (G) Boxplot showing performance score, stratified by regression-based procedures.

After quality control, we obtained an average about 49.2 million high-quality and uniquely mapped sequencing read pairs from each library (**Supplementary Fig. 1A**). Assessment of datasets corroborated that sequencing saturation has been reached (**Supplementary Fig. 1B**). Spike-in 2, which contained 5% NAD-capped forms, were significantly enriched, whereas no enrichment was found for spike-in 3 made up with 100% m^7^G-RNA (**Supplementary Fig. 1C**). Above evidence highlighted the sensitivity and specificity of the enrichment experiment, as reflected by the enrichment of NAD, but not m^7^G, capped transcripts.

### The feasibility of enONE

Since all samples were added with equal amounts of *Drosophila* spike-in RNAs, its disconcordance, if present, can be used to pinpoint the nuisance technical variation in an epitranscriptome-wide manner, and its concordance, on the other hand, can be used to validate the effect of normalization. To capture unwanted variation, i.e., batch effect, we use a set of genes (n = 1,000) whose expression patterns should be highly reproducible and now become differed among batches as the anchor set (**Fig. 2B**). In addition, we showed that normalization procedures were robust when the enrichment effect accounted for a small fraction of the anchor set variance, e.g., anchor set size ranged from 500 to 2,500 (**Fig. 2C and 2D**). With anchor set from *Drosophila* spike-in RNAs, we implemented enONE normalization procedures with five scaling toolsets, including TC, UQ, TMM, DESeq and PoissonSeq^17^, as well as three regression-based procedures, namely RUVg, RUVs, and RUVse. By integrating two normalization modules, we generated a total of 96 combinatorial procedures for the current data (**Fig. 2E**). By inspecting the full space of normalization performance metrics, we found that the top-ranked procedure involved DESeq scaling followed by RUVg adjustment for the first 4 factors of unwanted variation (**Fig. 2F and 2G**).

To validate the effect of enONE normalization, we applied RUVg (k = 4), RADAR, and the enONE procedure on all three types of spike-in RNAs. Compared to other procedures, enONE normalization dramatically mitigated the batch effect of *Drosophila* spike-ins in both input and enrichment libraries, while preserving the enrichment signals (**Fig. 3A**). In *Drosophila* spike-ins, linear regression between the first six PCs cumulatively and batch effect showed that enONE removed the variation among batches compared to other methods (**Fig. 3B**). Analysis of correlation between the gene normalized counts from *Drosophila* spike-ins and batch variation revealed a large proportion of genes showing strong correlations with batch effect in raw and RADAR normalized datasets, whereas this correlation was mitigated in the enONE normalized dataset (**Fig. 3C**). ANOVA was performed on *Drosophila* spike-ins from different batches, demonstrating the number of genes affected by batch effect was significantly reduced by enONE (**Fig. 3D**). Additionally, enONE improved the concordance of synthetic spike-ins compared to other methods (**Fig. 3E**). By employing the top-ranked normalization procedure on the analysis of PBMCs dataset, we noted that batch effect was mitigated while the enrichment covariates were well-preserved as illustrated by the first two principal components of PCA (**Supplementary Fig. 1D**). Together, these results demonstrated the capacity of enONE in removing unwanted variation while retaining the covariates of interest.

**Figure 3:**
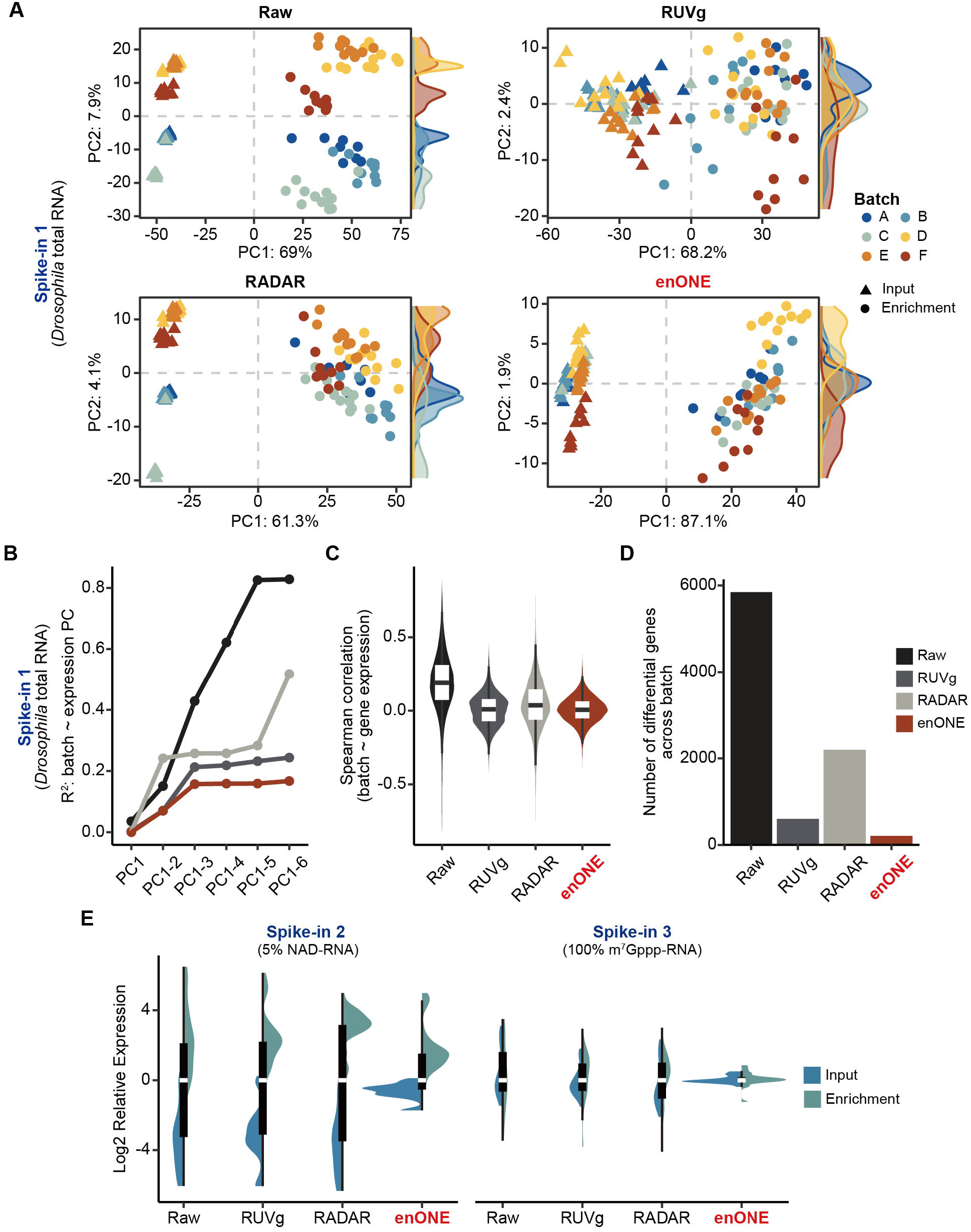
Performance assessment of different normalizations on spike-in controls. (A) PCA of *Drosophila* spike-in counts from different procedures. (B) The *R*^*2*^ of linear regression between batch effect and up to the first six PCs (taken cumulatively). (C) Spearman correlation coefficients between the normalized counts and batch effect. (D) The number of genes varied among bathes as inferred by ANOVA (*P* < 0.01). (E) Boxplot showing the relative expression levels of synthetic spike-ins with 5% NAD-caps and that with 100% m^7^G-caps from different methods.

### Characterization of NAD-RNAs from human PBMCs

We proceeded to set 2-fold enrichment of read counts as the cutoff, which led us to identify a total of 782 NAD-RNAs from human PBMCs (**Fig. 4A and Supplementary Table 2-4**). We then characterized these newly identified NAD-RNAs. In human PBMCs, NAD-capping mostly occurred on protein-encoding genes, but also extended to pseudogenes and non-coding RNAs, including lincRNA, snRNA, snoRNA, and miscRNA (**Fig. 4A**). NAD-RNAs were shown to be derived from genes localized on autosomes and X chromosomes, but not from the Y chromosome and the mitochondrion genome (**Fig. 4B**). By dividing NAD-RNAs into 5 deciles based on enrichment, we observed that shorter genes and genes with fewer introns tended to have increased modification of NAD (**Fig. 4C**), a pattern consistent with our recent study in mouse livers^8^. To inspect NAD modification of genes associated with biological functions, we performed pathway enrichment analysis, which revealed that NAD-RNAs were mainly involved in RNA metabolism, translation, transcription, energy metabolism, and immune system (**Fig. 4D and Supplementary Table 5**).

**Figure 4:**
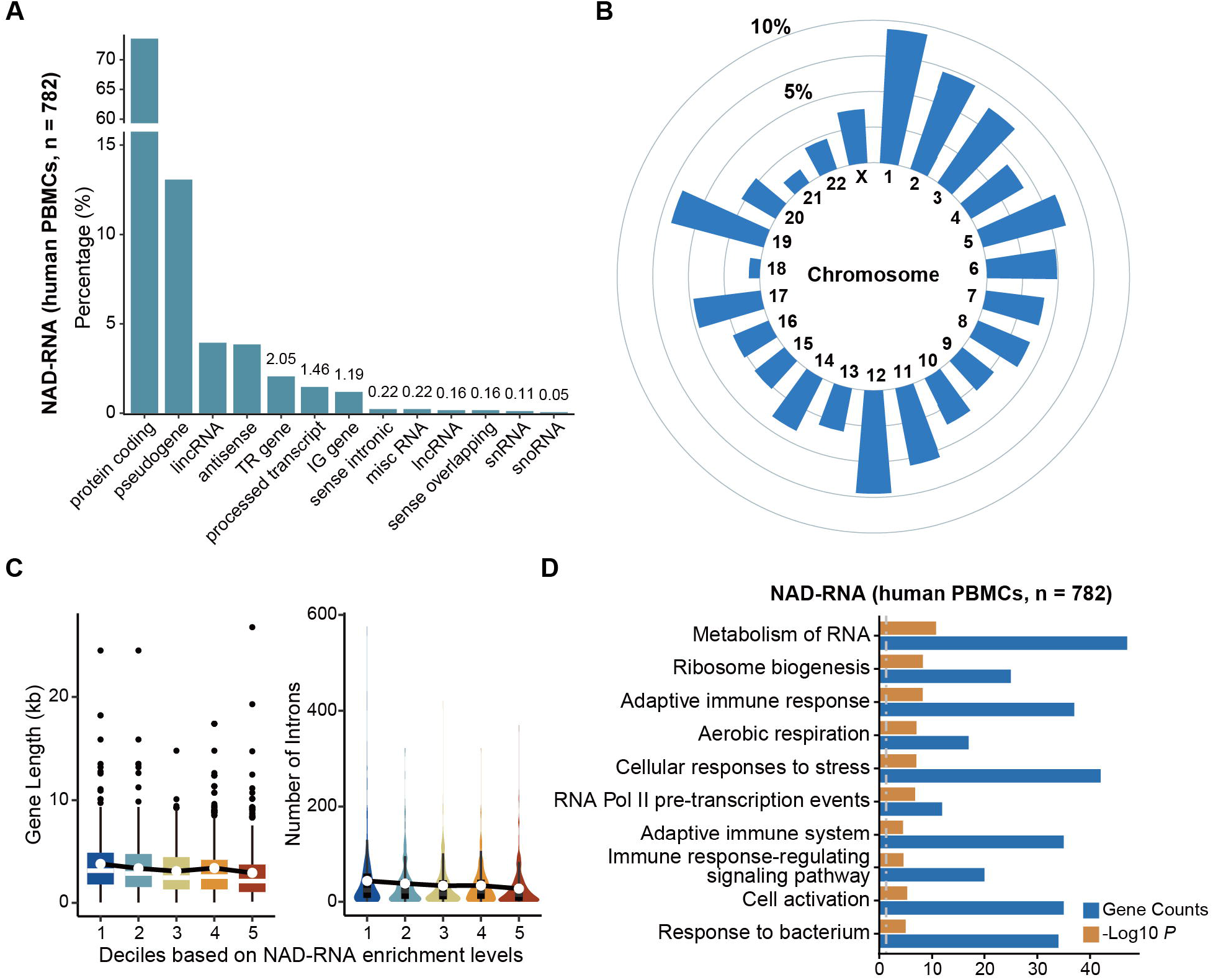
enONE facilitates NAD-capped RNAs identification. (A) Barplot showing the gene types of identified NAD-RNAs (n = 782) from human PBMCs. (B) Circular bar plot showing the chromosomes distribution of identified NAD-RNAs. (C) From five deciles based on enrichment, genes with short length and with fewer introns tend to have increased modification of NAD. (D) Pathway analysis reveals the biological processes of NAD-capped RNAs that are mainly involved in RNA metabolism, transcription, translation, and immune system. Grey dashed line in the bar plot indicates the 0.05 *P*-value cutoff.

### Age alters NAD-modified epitranscriptome

To gain insights into how NAD-RNAs are modulated with age and its consequent impact on the progression of aging, we analyzed NAD-RNA profiles from all age groups. Interestingly, despite the fact that NAD decreases with age^18^, we found that the number of NAD-capping events tended to increase in aged human subjects (**Fig. 5A**). To dissect this observation, we grouped age-associated trajectories into three major clusters using hierarchical clustering (**Fig. 5B and Supplementary Table 6**). Increased NAD modification was found for genes in cluster 1, with their function being involved in basic cellular events and adaptive immune response. In cluster 2, NAD-capping was increased in early age but later became plateaued; these genes were functionally enriched in oxidative stress and innate immune response. Genes associated with collagen production, protein phosphorylation, and TGF-β signaling pathway, which were ascribed as cluster 3, exhibited a decreased trend in NAD modification (**Fig. 5B and Supplementary Table 7**). Further analyses revealed that genes in cluster 3, with their expression and NAD-modification, were well-correlated, whereas genes in cluster 1 and 2 were less-correlated (**Supplementary Fig. 2A and 2B**).

**Figure 5:**
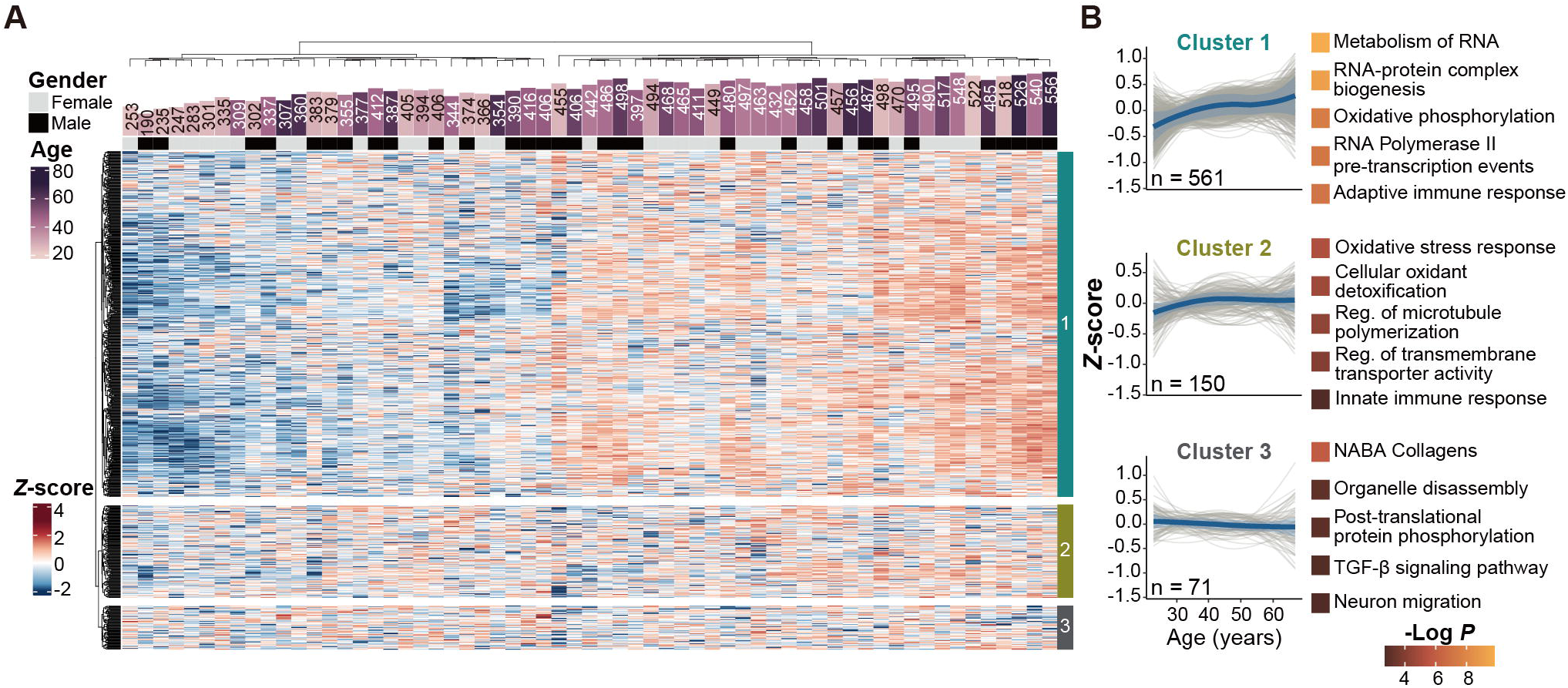
enONE reveals dynamics of NAD-capped RNAs during aging. (A) Heatmap showing epitranscriptomic profiles from individuals across age groups (N = 61). Top bar represents the enrichment signals (Fold-Change ≥ 2) in each sample. (B) Unsupervised hierarchical clustering was used to group NAD-RNAs with similar trajectory. Three major clusters were identified and the top five non-redundant pathways were listed. The solid line and shaded region represent the smoothed trajectory and its 95% confidence intervals, respectively.

By inspecting the correlation between NAD modification and age, we identified a set of NAD-RNAs that highly associated with age (n = 67) (**Fig. 6A**). Specifically, select NAD-RNAs, such as those involved in protein folding (PDIA3), protein ubiquitination (SUMO1), and apoptosis (caspase 3 and 8), had increased capping with age (**Supplementary Table 6**); but the abundance at RNA transcript levels was not increased (**Fig. 6B**). In addition, NAD-capping of genes linked to mRNA decay (UPF2), calmodulin binding (NRGN), and TGF-β signaling pathway (TGFB1) were decreased during aging (**Supplementary Table 6**). The expression levels of UPF2 and NRGN also decreased during aging, while TGFB1 was increased (**Fig. 6B**). Together, our study revealed the first NAD-modified epitranscriptome from human PBMCs. Using enONE, we were able to pinpoint epitranscriptomic alteration of NAD-capping during the aging process.

**Figure 6:**
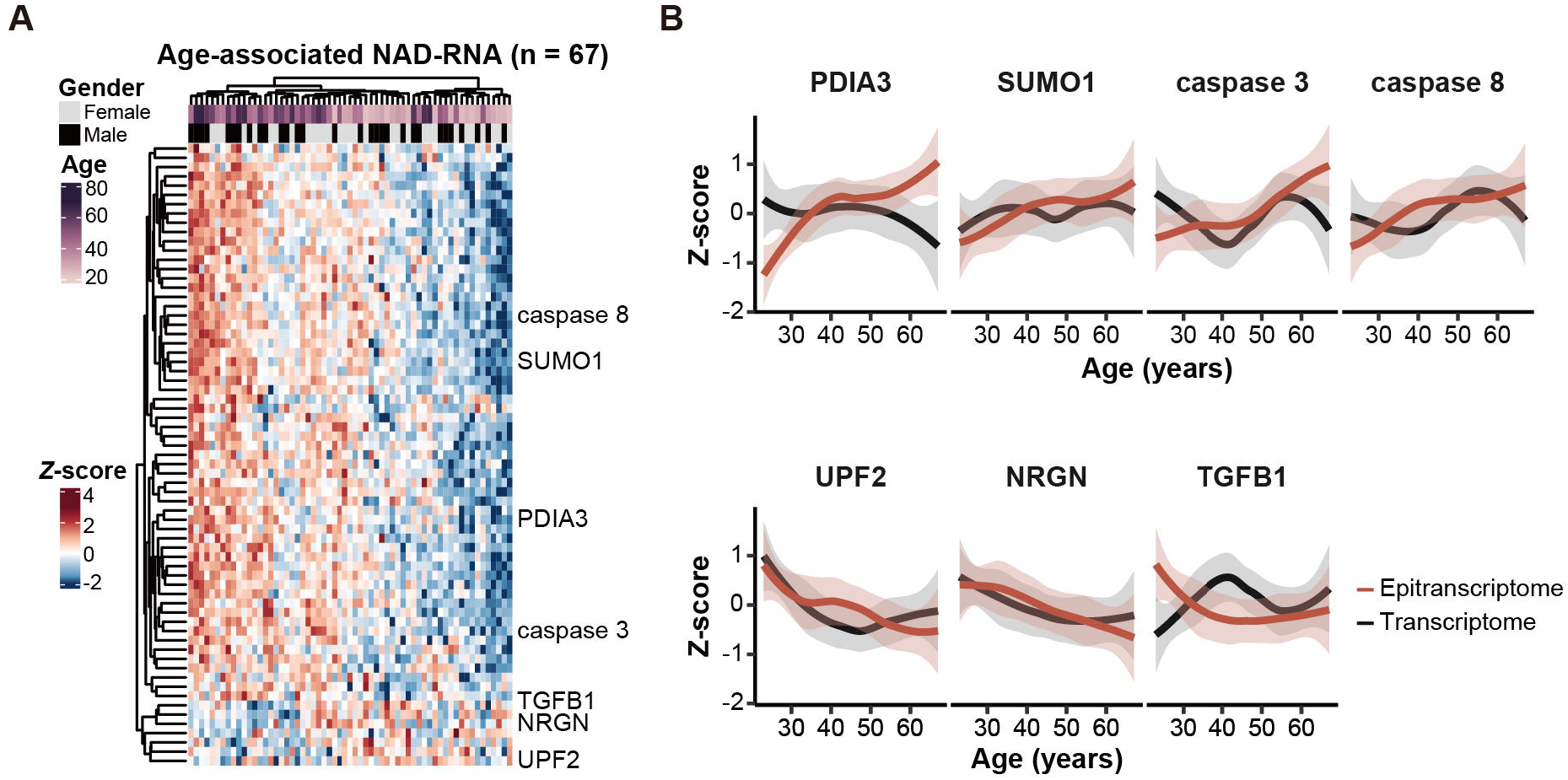
NAD-capped RNAs reflect the processes of aging. (A) Heatmap showing a set of NAD-RNAs (n = 67) that highly associated with age. (B) *Z*-transformed, smoothed trajectories of NAD-capping levels (in red) and expression levels (in black) of selected genes were shown. The solid line and shaded region represent the smoothed trajectory and its 95% confidence intervals, respectively.

## DISCUSSION

Incorporation of NAD, the hub metabolite and redox cofactor for cells, into RNA represents the crosstalk between metabolite and gene expression, defining a critical layer of epitranscriptomic regulation. The recently developed NAD-RNA sequencing technologies have substantially facilitated the epitranscriptome-wide identification of novel NAD-capped RNA across phyla^19^. However, since these methods require affinity-based enrichment, data normalizations borrowed from bulk RNA-seq could not be readily applied. In this study, we devise a general framework for spike-in-based epitranscriptome-wide NAD-RNA-seq data normalization and evaluation. Compared to previous epitranscriptome profiling with ERCC spike-ins, enONE includes *Drosophila* total RNA as spike-in to account for the effect of affinity-based enrichment in an epitranscriptome-wide manner. Scaling-based approaches have been extensively used in the analysis of epitranscriptome profiles, whereas we demonstrate that integration of global scaling and RUV strategies can optimize normalization performance (see **Fig. 2G**). Selection of the “best” normalization, however, might not be feasible in practice due to the subjective definition of optimality. Therefore, enONE emphasizes the choice of an appropriate strategy rather than the “best” normalization. Together, enONE is able to efficiently remove the impact of unwanted variations caused by affinity-based enrichment from the data. We highlight the application that enONE, an open-source R package, can be extended into the analysis of other types of epitranscriptomic sequencing data that involve step-wise enrichment procedures, e.g., m^6^A-seq.

enONE facilitates the identification of NAD-RNAs. As a source of liquid biopsy, peripheral blood can serve as sentinel tissue to monitor individual health in a non-invasive manner. Identification of novel blood-derived features may open up new avenues in detection of biomarkers for various physiological and pathological conditions. Here we reveal prominent features of NAD-RNAs from human PBMCs. Large collections of NAD-RNAs are produced by protein-encoding genes, with their biological functions mainly involved in basic cellular events, such as translation, RNA metabolism, and transcription. Additionally, cell-specific NAD-RNAs are discovered, such as those involved in immune system. Thus far, yet the function of NAD-RNAs remains elusive, their dynamic changes may provide insights into how individuals respond to perturbations.

We reveal the dynamics of NAD-modified epitranscriptome during aging. Though numerous studies have shown that NAD metabolite decreases with age^16,18^, our data demonstrate increased NAD-capping in elder population, suggesting that the addition or removal of 5’-terminus NAD moiety might not be solely dependent on the cellular reserve of NAD (see **Fig. 5A**). Notably, some of the age-associated NAD-RNAs are involved in molecular pathways profoundly impinging on the hallmarks of aging, such as ribosome biogenesis, immune system, and mitochondrial function^20^, thus defining a novel mechanistic component of the aging process. Yet, we acknowledge the limitation. Our human aging cohort is from the local community of Shanghai, which does not represent the general population. Intriguingly, the overall number of NAD-RNAs from individual subjects is not simply correlated with age (see **Fig. 5A**), raising the possibility that additional age-associated factors might be linked. Since there is currently no gold standard measure of biological aging^21^, it is tempting to exploit how NAD-capped RNAs as biological implications might signify physiological and perhaps pathological conditions.

Taken together, we propose enONE as a flexible, modular, and general framework for the normalization and evaluation of NAD-RNA sequencing data in a wide spectrum of biological contexts. To the best of our knowledge, enONE is the first computational approach for NAD-modified epitranscriptome profiles. Future characterization of NAD-capped RNAs, empowered by enONE, can focus on how biological processes regulate NAD modification through a quantitative lens, especially during the aging process.

## Methods

### Ethics Statement

The project was approved by the Medical Ethics Committee of Shanghai Changzheng Hospital (2022SL006). Informed consent was obtained from all subjects in accordance with the local research Ethics Committee guidelines.

### Study design and participants

We conducted a cross-sectional study of natural aging in community subjects recruited from Shanghai Changzheng Hospital. The aging cohort consists of 31 females and 30 males, aged from 23 to 67 years old. Inclusion criteria for cohort were: 1) above 20 years old; 2) independently able to provide written informed consent. For comparative analysis of age-related subgroups, the cohort was divided into three age groups: Young (20-35 years old), Middle (36-50 years old), and Old (51 years old and above). Height and weight are measured by trained staff following standardized protocols. BMI (kg/m^2^) is derived from the calculation and stored for further analyses. After 5 minutes of rest in the seated position, blood pressure (mmHg) was measured three times with an automatic sphygmomanometer and the mean of the measurements was used for analysis. Blood samples were taken from all patients in the morning after they had been seated for 5 minutes. All participants had blood drawn using lithium heparin tubes (BD Vacutainer, catalog: 367884) by phlebotomists and consented to having their de-identified survey data made publicly available.

### Isolation of peripheral blood mononuclear cells (PBMCs) from whole blood

For isolation of peripheral blood mononuclear cells (PBMCs), 5 mL whole blood was mixed with 630 μL OptiPrep (Sigma-Aldrich, catalog: D1556) and 500 μL solution C (0.85% (w/x) NaCl and 10 mM Tris-HCl, pH 7.4), followed by centrifugation at 4 °C and 1,300 g for 30 min. PBMCs were collected and mixed with two volumes of solution C, followed by centrifugation at 4 °C and 500 g for 10 min. Cell pellet was washed twice with 1 mL PBS. The suspension was used for NAD-RNA detection.

### In vitro transcription of NAD-RNA, and m^7^Gppp-RNA

To assess the sensitivity of enrichment procedures, spike-in NAD-RNA (500 nt; sequence A) and m^7^Gppp-RNA (500 nt; sequence A) with identical sequence, oligonucleotide without adenine was synthesized (Genewiz) and were subjected to polyadenylation for poly(A) tails elongation (template sequence: 5′-**TAATACGACTCACTATT**ATGGTGTGCTTGGGCGTGGTGCTGTTCTCCGGGG TGGTGCCCTTCCTGGTCGTGCTGGTCGGCGTCGTTTTCGGCCTCTTGTTCTGCGTGTCCGGCGTGGGCGTGGGCGTTGCCTCCTTCGGCTTGCTGTCCCTGTTGTTCTTCTGCTCCTCCGGCTTGCTGCCCGTGCCCTGGCCCTCCCTCGTGTCCTCCCTGTCCTTCGGCGTGCTGTGCTTCTGCCGCTTCCCCGTCCTCTTGTTGCTGCTCGTCTTCTTCTTGTCCGCCTTGCCCGTTGGCTTCGTCCTGGTGCGCTCCTTCTTCTTCTTGGTCGTCGGCTTCTTCTTGTCCCGCGCCGTGGTGTTGTTC GTGGGCGTCTCCCTGGTGTTCCGCTTCGTGCTGTTGGGCTTCGTCTTCTTGGTGGTCGGCTTCTTCCTGGGGCTCTTGCTGGTGTTCTTCTTCTTCTGCCTCTTCGTCTTTTTCTTGGCCGTCTTGCTGTTGTTCGGCTTCTTGGTGTTCTTCTT-3′; boldface letters denote the sequence of T7 class II promotor (□2.5)) and (anti-sense: 5′-AAGAAGAACACCAAGAAGCCGAACAACAGCAAGACGGCCAAGAAAAAGACGAAGAGGCAGAAGAAGAAGAACACCAGCAAGAGCCCCAGGAAGAAGCCGACCACCAAGAAGACGAAGCCCAACAGCACGAAGCGGAACACCAGGGAGACGCCCACGAACAACACCACGGCGCGGGACAAGAAGAAGCCGACGACCAAGAAGAAGAAGGAGCGCACCAGGACGAAGCCAACGGGCAAGGCGGACAAGAAGAAGACGAGCAGCAACAAGAGGACGGGGAAGCGGCAGAAGCACAGCACGCCGAAGGACAGGGAGGACACGAGGGAGGGCCAGGGCACGGGCAGCAAGCCGGAGGAGCAGAAGAACAACAGGGACAGCAAGCCGAAGGAGGCAACGCCCACGCCCACGCCGGACACGCAGAACAAGAGGCCGAAAACGACGCCGACCAGCACGACCAGGAAGGGCACCACCCCGGAGAACAGCACCACGCCCAAGCACACCATAATAGTGAGTCGTATTA-3′). To assess the specificity of enrichment procedures, spike-in m^7^Gppp-RNA (500 nt; sequence B) oligonucleotide was synthesized (Genewiz) and were subjected to polyadenylation for poly(A) tails elongation (template sequence: 5′-**TAATACGACTCACTATT**ACATGGAGGGCTCCGTGAACGGCCACGAGTTCG AGATCGAGGGCGAGGGCGAGGGCCGCCCCTACGAGGGCACCCAGACCGCCAAGCTGAAGGTGACCAAGGGTGGCCCCCTGCCCTTCGCCTGGGACATCCTGTCCCCTCAGTTCATGTACGGCTCCAAGGCCTACGTGAAGCACCCCGCCGACATCCCCGACTACTTGAAGCTGTCCTTCCCCGAGGGCTTCAAGTGGGAGCGCGTGATGAACTTCGAGGACGGCGGCGTGGTGACCGTGACCCAGGACTCCTCCCTGCAGGACGGCGAGTTCATCTACAAGGTGAAGCTGCGCGGCACCAACTTCCCCTCCGACGGCCCCGTAATGCAGAAGAAGACCATGGGCTGGGAGGCCTCCTCCGAGCGGATGTACCCCGAGGACGGCGCCCTGAAGGGCGAGATCAAGCAGAGGCTGAAGCTGAAGGACGGCGGCCACTACGACGCTGAGGTCAAGACCACCTACAAGGCCA-3′; boldface letters denote the sequence of T7 class II promotor (□2.5)) and (anti-sense: 5′-TGGCCTTGTAGGTGGTCTTGACCTCAGCGTCGTAGTGGCCGCCGTCCTTCA GCTTCAGCCTCTGCTTGATCTCGCCCTTCAGGGCGCCGTCCTCGGGGTACA TCCGCTCGGAGGAGGCCTCCCAGCCCATGGTCTTCTTCTGCATTACGGGGC CGTCGGAGGGGAAGTTGGTGCCGCGCAGCTTCACCTTGTAGAT GAACTCGCCGTCCTGCAGGGAGGAGTCCTGGGTCACGGTCACCACGCCGC CGTCCTCGAAGTTCATCACGCGCTCCCACTTGAAGCCCTCGGGGAAGGAC AGCTTCAAGTAGTCGGGGATGTCGGCGGGGTGCTTCACGTAGGCCTTGGA GCCGTACATGAACTGAGGGGACAGGATGTCCCAGGCGAAGGGCAGGGGG CCACCCTTGGTCACCTTCAGCTTGGCGGTCTGGGTGCCCTCGTAGGGGCGG CCCTCGCCCTCGCCCTCGATCTCGAACTCGTGGCCGTTCACGGAGCCCTCC ATGTAATAGTGAGTCGTATTA-3′). For *in vitro* transcription, 10 μM of double-stranded DNA (dsDNA) template in 100 μL transcription buffer (Promega, catalog: P1300), along with 1 mM of each of GTP, CTP and UTP, with 4 mM NAD (for NAD-RNA) or 4 mM m^7^GpppA (New England Biolabs, catalog: S1406S) (for m^7^G-RNA), 10 μL of T7 RNA polymerase (Promega, catalog: P1300), 5% DMSO, 5 mM DTT and 2.5-unit RNase inhibitor were added and the transcription mixture was incubated at 37 °C for 4 h. The reaction was incubated with 11-unit DNase I (Promega, catalog: P1300) at 37 °C for 30 min to remove the DNA template. RNA was then extracted using acid phenol/chloroform and precipitated with isopropanol (with 0.3 M sodium acetate, pH 5.5) at -80 °C overnight. The RNA pellet was washed twice with 75% ethanol, air-dried, re-dissolved in DEPC-treated H_2_O, and stored at -80 °C.

### NAD-capped RNA sequencing

Total RNAs from human PBMCs and total RNAs from *Drosophila* (spike-in) were prepared in accordance with the manufacturer’s instruction (Takara Bio, catalog: 9108). Total RNAs (10 μg) from human PBMCs were mixed with 40 μg *Drosophila* RNA (spike-in 1), 0.1 ng synthetic RNAs (spike-in 2: 5% NAD-RNA/95% m^7^G-RNA; sequence A), and 0.1 ng synthetic RNAs (spike-in 3: 100% m^7^G-RNA; sequence B). The mixture of total RNAs and spike-in RNAs was incubated with 100 mM HEEB (1 M stock in DMSO) with ADPRC (25 μg/mL) in 100 μL of ADPRC reaction buffer (50 mM Na-HEPES pH 7.0, 5 mM MgCl_2_) at 37 °C for 1 h, followed by NudC-catalyzed NAD-RNA elution. 100 μL of DEPC-treated H_2_O was then added and acid phenol/ether extraction was performed to stop the reaction. RNAs were precipitated by ethanol, and re-dissolved in 100 μL of DEPC-treated H_2_O. 5 μL of biotinylated RNAs were kept as input. After HEEB reaction, biotinylated RNAs were incubated with streptavidin bead particles (6 μL, MedChemExpress, catalog: HY-K0208) and 0.4 U/μL of RNase Inhibitor (Takara Bio, catalog: 2313B) at 25 °C for 30 min. Beads were washed four times with streptavidin wash buffer (50 mM Tris-HCl (pH 7.4) and 8 M urea), and three times with DEPC-treated H_2_O. To ensure complete elution, biotin-conjugated RNAs were replaced from streptavidin beads by incubating with 1 mM biotin buffer (20 μL, Sigma-Aldrich, catalog: B4639) at 94 °C for 8 min, followed by incubation with 500 nM NudC (New England Biolabs, catalog: M0607S) in 25 μL of NudC reaction buffer (100 mM NaCl, 50 mM Tris-HCl pH 7.9, 10 mM MgCl_2_, 100 µg/ml Recombinant Albumin) at 37 °C for 30 min. After NudC treatment, biotinylated-RNAs that are resistant to NudC catalysis, potentially derived from contaminating m^7^G-RNAs, were retained on beads by incubation with high-capacity streptavidin particle (20 μL, Thermo Fisher Scientific, catalog: 20357) at 25 °C for 30 min. Eluted RNAs in the supernatant were used for next step. Input (see above) and NudC-eluted RNAs were used for NGS library construction, in accordance with the manufacturer’s instructions (mRNA-seq Lib Prep Kit for Illumina, Abclonal, catalog: RK20302). Library quality was assessed by Bioanalyzer 2100 (Agilent, United States), and quantification was performed by qRT-PCR with a reference to a standard library. Libraries were pooled together in equimolar amounts to a final 2 nM concentration and denatured with 0.1 M NaOH (Sigma, catalog: 72068). Libraries were sequenced on the Illumina NovaSeq 6000 system (paired end; 150 bp).

### High-throughput sequencing data analysis

All sequencing reads were processed with Trim Galore (v0.6.6)^22^ with the parameters “--nextseq 30 --paired” to remove the adapter sequences (AGATCGGAAGAGC) from NovaSeq-platforms, and reads longer than 20 bp were kept. Reads that passed the quality control procedure were kept and mapped to the *Homo sapiens* genome (GRCh38) and *Drosophila melanogaster* genome (dmel-all-chromosome-r6.36) using STAR (v2.7.6a)^23^ with default parameters, respectively. Uniquely mapped read pairs were counted against annotations from *Homo sapiens* (Ensembl: GRCh38.94) and *Drosophila melanogaster* (Flybase: dmel-all-r6.36) using featureCounts (v2.0.1)^24^ with parameters “-p -B -C” and summarized as gene-level counts, respectively. Sequencing saturation was assessed by randomly subsampling the original libraries and examined the corresponding changes in the number of genes, derived from human genome, with more than 10 read counts.

### enONE workflow

enONE is implemented in R and publicly available at https://github.com/thereallda/enONE. enONE workflow consists of four steps: 1. Quality control; 2. Gene set selection; 3. Normalization procedures; 4. Normalization performance assessment. By “log” transformation, we generally refer to the log_2_(x+1) function unless otherwise stated. Below, steps are shown in details.

#### 1. Quality control

The goal of quality control was to remove problematic or noisy observations from downstream analysis. In this study, we used sample and gene filtering to control data quality. To assess outliers, we applied Rosner’s outlier test on principal component 1. Principle component analysis (PCA) was performed with prcomp function on the top 20,000 genes based on a transformed counts matrix by vst function from R package DESeq2 (v1.36.0)^13^. All samples were kept for subsequent analysis. To keep well-detected genes across samples, we used filterByExpr function from the R package edgeR (v3.38.4)^25^ with parameter “min.count=20”. All ribosomal RNA encoded genes and TEC genes were excluded.

#### 2. Gene set selection

enONE defined three sets of control genes for adjustment of the unwanted variations, evaluation of the unwanted variations, and evaluation of the wanted variations, respectively. For adjustment of the unwanted variation, we defined the 1,000 least significantly enriched genes in *Drosophila* spike-ins, ranked by FDR values, as negative control genes. Since the effect of affinity-based enrichment was the known covariate of interest, we used these genes, that were not affected by the enrichment effect, to compute the unwanted variation in the subsequent RUV procedure. For evaluation of the unwanted variation, we defined the 500 least significantly varied genes in human, ranked by FDR values, as negative evaluation genes. We performed differential analysis test across all covariates of interest to determine genes with constant expression levels. Since the variation of constant genes could reflect the handling effects, those genes could be used to evaluate the removal of unwanted variation in the subsequent normalization evaluation step. For evaluation of the wanted variation, we defined the 500 most significantly enriched genes in human, ranked by FDR values, as positive evaluation genes. We performed differential analysis test between enrichment and input samples in human to determine genes affected by enrichment procedures. Those genes were used to evaluate the preservation of wanted variation in the subsequent normalization evaluation step.

#### 3. Normalization procedures

enONE implemented global scaling and regression-based methods for the generation of normalization procedures. For the global scaling normalization procedures, gene-level read counts were scaled by a sample-wise scale factor. Five different scaling procedures were implemented, including: 1) Total Count (TC): The scale factor was defined as the sum of the read counts across all genes. 2) Upper-Quartile (UQ): The scale factor was defined as the upper-quartile of the gene-level count distribution. 3) TMM^12^: The scale factor was based on a robust estimate of the overall expression fold change between the sample and a reference sample. TMM was performed by the calcNormFactors function from R package edgeR with parameter: method = “TMM”. The default behavior used here is that the selected reference sample has an upper quartile closest to the mean upper quartile of all samples. 4) DESeq^13^: The scale factor for a given sample was defined as the median fold-change between the samples and a pseudo-reference sample whose counts were defined as the geometric means of the counts across samples. This method was performed by the calcNormFactors function from R package edgeR with parameter: method = “RLE”. Note that the method discarded any gene having zero count in at least one sample. 5) PoissonSeq: The scale factor was based on an iterative estimate of sequencing depth from a subset of genes, defined by a Poisson goodness-of-fit statistics^17^. By default, this method discarded genes with less than 5 read counts.

For the regression-based procedures, enONE considered the following generalized linear model (GLM), which allows adjustment for factors of unwanted variation:

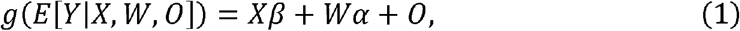

where *Y* was the *n* x *J* matrix of gene-level read counts, *X* was an *n* x *M* design matrix corresponding to the *M* covariates of interest of ‘‘wanted variation’’ (e.g., treatment) and β its associated *M* x *J* matrix of parameters of interest, *W* was an *n* x *K* matrix corresponding to unknown factors of unwanted variation and α its associated *K* x *J* matrix of nuisance parameters, *O* was an *n* x *J* matrix of offsets that can either be set to zero or estimated with other normalization procedure (such as TMM normalization), and *g* was a link function.

The unwanted variation *W* can be estimated by singular value decomposition (SVD) using main approaches as implemented in R package RUVSeq (v1.30.0)^14^. In this study, we used three variants of RUV, including: 1) RUVg: estimated the factors of unwanted variation based on negative control genes, assumed to have constant expression across all samples (β=0), as applied in RUV. 2) RUVs: estimated the factors of unwanted variation based on negative control genes from the replicate samples (e.g., replicates in each treatment group) for which the covariates of interest were constant (β=0), as applied in RUV. 3) RUVse: was a modification of RUVs. It estimated the factors of unwanted variation based on negative control genes from the replicate samples in each assay group (i.e., enrichment and input), for which the enrichment effect was assumed to be constant.

#### 4. Normalization performance evaluation

To evaluate the performance of normalization, we leveraged eight normalization performance metrics that related to different aspects of gene expression measures^26^.

The following four metrics evaluated normalization procedures based on how well the samples can be clustered according to factors of wanted and unwanted variation. Clustering by wanted factors was desirable, while clustering by unwanted factors was undesirable. As clustering measurements, we used silhouette widths. For any clustering of *n* samples, the silhouette width of sample *i* was defined as:

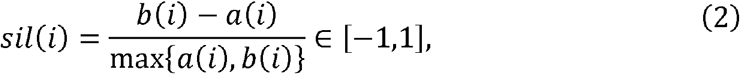

where *a(i)* denoted the average distance (by default, Euclidean distance over the first three PCs of expression measures) between the *i*th sample and all other samples in the cluster to which *i* was assigned and *b(i)* denoted the minimum average distance between the *i*th sample and samples in other clusters. The larger the silhouette widths, the better the clustering. Thus, the average silhouette width across all *n* samples provides an overall quality measure for a given clustering. Silhouette width was calculated with the silhouette function from R package cluster (v2.1.3). These metrics were defined as follows: 1) BIO_SIM: Group the *n* samples according to the value of a categorical covariate of interest (e.g., age group or treatment) and compute the average silhouette width for the resulting clustering. 2) BATCH_SIM: Group the *n* samples according to the batch and compute the average silhouette width for the resulting clustering. 3) EN_SIM: Group the *n* samples according to the assay (e.g., enrichment or input) and compute the average silhouette width for the resulting clustering. 4) PAM_SIM: Cluster the *n* samples using partitioning around medoids (PAM) for a range of numbers of clusters and compute the maximum average silhouette width for these clusters. PAM clustering was done by the pamk function from R package fpc (v2.2-9). Large values of BIO_SIM, EN_SIM and PAM_SIM and low values of BATCH_SIM were desirable.

The next two metrics evaluated the association of log-count principal components (by default, the first 3 PCs) with “evaluation” principal components of wanted or unwanted variation. 1) UV_COR: The weighted coefficient of determination 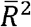 for the regression of log-count principal components on factors of unwanted variation (UV) derived from negative evaluation genes. The submatrix of log-transformed unnormalized counts for negative evaluation genes is row-centered and scaled and factors of unwanted variation are defined as the right-singular vectors as computed by the svd function. 2) WV_COR: The weighted coefficient of determination 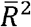 for the regression of log-count principal components on factors of wanted variation (WV) derived from positive evaluation genes. The WV factors were computed in the same way as the UV factors above, but with positive instead of negative evaluation genes. Large values of WV_COR and low values of UV_COR were desirable.

The weighted coefficients of determination were computed as follows. For each type of evaluation criterion (i.e., UV, or WV), regressed each expression PC on all supplied evaluation PCs. Denoted *SST*_*k*_ as the total sum of squares, *SSR*_*k*_ as the regression sum of squares, and *SSE*_*k*_ as the residual sum of squares for the regression for the *k*th expression PC. The coefficient of determination was defined as:

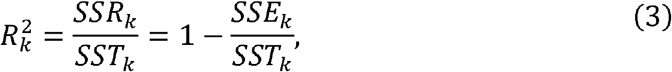

and weighted average coefficient of determination as:

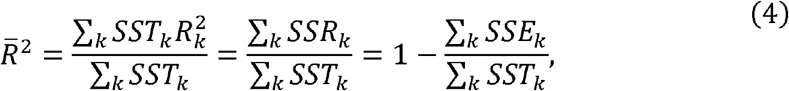

The next two metrics evaluated the similarity of gene expression distributions. We defined gene-level relative log-expression (RLE) measures, as log-ratios of read counts to median read counts across samples, for comparing distribution of gene expression. RLE was defined as follows:

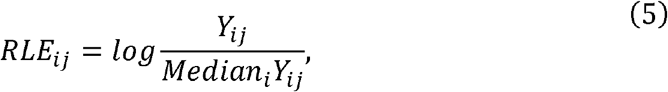

for gene *i* in sample *j*. For similar distributions, the RLE should be centered around zero and have a similar distribution across samples. Thus, the metrics were defined as follows: 1) RLE_MED: Mean squared median RLE. 2) RLE_IQR: Variance of inter-quartile range (IQR) of RLE. Low values of RLE_MED and RLE_IQR were desirable.

In the enONE framework, the expression measures were normalized according to a set of methods (including raw counts) and the eight metrics above were computed for each dataset. The performance assessment results can be visualized using biplots and the normalization procedures were ranked based on a function of the performance metrics. In particular, enONE defined a performance score by orienting the metrics (multiplying by ±1) so that large values correspond to good performance, ranking procedures by each metric, and averaging the ranks across metrics.

### Quantitative NAD-RNA-seq data analysis

To identify NAD-RNA from sequencing data, we performed differential analysis using FindEnrichment function from enONE package. Significance of logarithmic fold changes was determined by a likelihood ratio test to approximate *P* values, and genes were adjusted for multiple testing using the Benjamini-Hochberg procedure to yield a false discovery rate (FDR). NAD-RNAs were defined as fold change of normalized transcript counts ≥ 2, FDR < 0.05, and log_2_-CPM ≥ 1 in enrichment samples compared to those in input samples. The fold change of normalized counts between pairwise enrichment and input sample was denoted as NAD modification level. Gene annotation information, such as chromosome, gene types, and gene lengths were retrieved from Ensembl (GRCh38.94) annotations. The violin plot, boxplot, bar plot, line chart and scatter plot were generated by R package ggplot2 (v3.3.6)^27^.

### Pathway enrichment analysis

Pathway enrichment analysis was performed using Metascape (v3.5)^28^. Pathways were defined as the molecular pathways of Reactome, the biological process (BP) of GO, the Kyoto Encyclopedia of Genes and Genomes (KEGG), the WikiPathways, the canonical pathways of MSigDb, and the CORUM database. Metascape clustered enriched terms into non-redundant groups based on similarities between terms and used the most significant term within each cluster to represent the cluster. The resulting clusters of pathways were manually reviewed. Enrichment was tested using the hypergeometric test. Multiple hypothesis correction was performed with Benjamini-Hochberg procedure, and the significance threshold was defined as α = 0.05.

### Hierarchical clustering analysis

Hierarchical clustering with a Euclidean distance metric was performed for both rows (NAD-RNAs, complete method) and columns (samples, ward’s method) on the scaled NAD modification matrix. Clustering and corresponding heatmaps were generated by R package Complex Heatmap (v2.12.1)^29^. Using cutree function with “k=3” parameter, we identified 3 clusters of NAD-RNAs changing with age, ranging from 71 to 561 NAD-RNAs.

### Trajectory analysis

To estimate NAD modification trajectories during aging, log NAD modification levels were *z*-scored and LOESS regression was fitted for each gene. Similarly, log expression levels were used to estimate the gene expression trajectories during aging. The trajectory of clusters was estimated using the median levels of genes in each cluster. Pathways were queried as above to gain insights into the biological functions of each cluster. Top five non-redundant enriched terms were shown. To identify NAD-RNAs that correlated with age, we computed Spearman’s rank correlation coefficients (*r*_*s*_) for each NAD-RNAs on the basis of NAD modification and age. The resulting correlation metrics was then filtered, such that only NAD-RNAs that were significantly correlated with age (*P* < 0.05 and | *r*_*s*_ | ≥ 0.33) were considered. Similarly, correlation between age and gene expression levels was computed.

### Statistical analyses

All *P* values reported herein were calculated using the non-parametric Mann-Whitney rank test unless otherwise stated.

## DATA AVAILABILITY

All high-throughput RNA sequencing data as well as transcript quantifications have been deposited at the Gene Expression Omnibus under accession number GSE226636 (secure token for data review: mdctksagvhmndod). All source codes for data analysis are available at GitHub (https://github.com/thereallda/enONE-paper). enONE is available as an R package on GitHub (https://github.com/thereallda/enONE).

## ACKNOWLEDGEMENTS

Author contributions: Conceptualization, N.L. and C.Z.; Methodology, N.L., D.L.; Investigation, D.L., S.G., Y.L., K.N., M.P., X.W., G.H., S.Z., R.L., C.Z. and N.L.; Data Curation, D.L., and N.L.; Writing-Original Draft, D.L. M.P., C.Z. and N.L.; Writing-Review & Editing, D.L. M. P., C.Z. and N.L.; Funding Acquisition, L.Q., C.Z. and N.L.; Supervision, L.Q., C.Z. and N.L..

