## Supplementary material for "Epitranscriptome analysis of NAD-capped RNA by spike-in-based normalization": Suplementary Figures

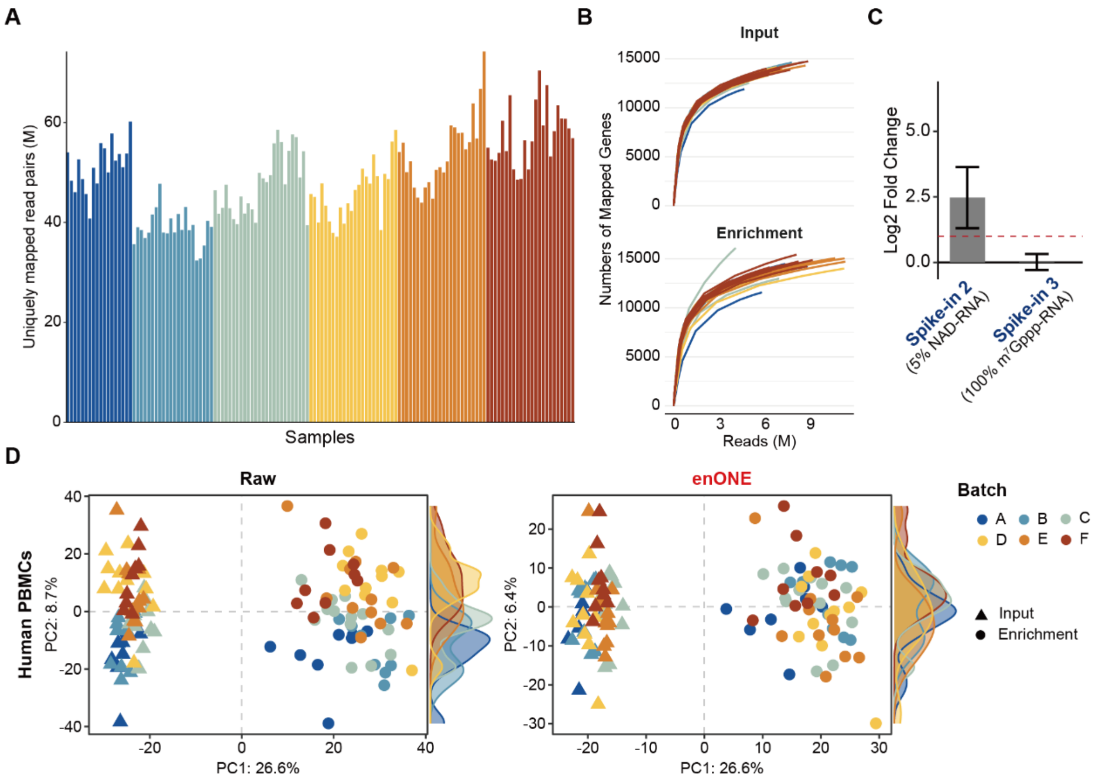


Li et al. Supplementary Figure 1

**Supplementary Figure 1: Quality assessment of NAD-RNA-seq data.** (A) Analysis of sequences alignment from human PBMCs. (B) Analysis of sequencing saturation for the human genome. (C) Barplot showing the fold change between enrichment and input samples of synthetic spike-ins with 5% NAD-caps and that with 100% $m^7$G-caps. Red dashed line represents the 2-fold enrichment cutoff. Data are mean ± se. (D) Analysis of PCA for human profiles with or without top-ranked normalization.

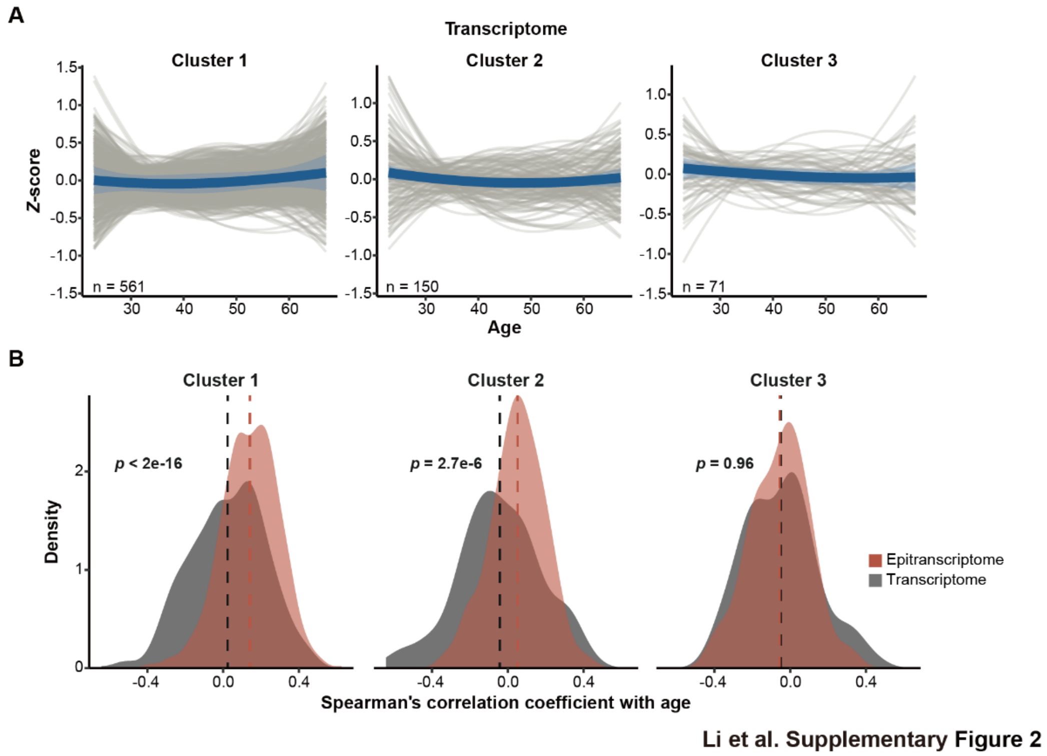


**Supplementary Figure 2: Assessment of Transcriptome of each cluster.** (A) Gene expression trajectories of each cluster based on NAD-capping dynamics. The solid line and shaded region represent the smoothed trajectory of each cluster and its 95% confidence intervals, respectively. (B) Except for cluster 3, correlation between age and NAD modification differed significantly from that between age and gene expression (*p*-values were assessed using the Kolmogorov-Smirnov test). Dashed lines represent the average correlation of each distribution.
